# Adoptive transfer of CD49a^+^ Tissue resident memory cells reverses pulmonary fibrosis in mice

**DOI:** 10.1101/2024.03.13.584814

**Authors:** Samuel L Collins, Yee Chan-Li, Kevin Shenderov, Astrid Gillich, Andrew M. Nelson, Jeffrey M. Loube, Wayne A Mitzner, Jonathan D. Powell, Maureen R. Horton

## Abstract

Pulmonary fibrosis is a devastating disease with no effective treatments to cure, stop or reverse the unremitting, fatal fibrosis. A critical barrier to treating this disease is the lack of understanding of the pathways leading to fibrosis as well as those regulating the resolution of fibrosis. Fibrosis is the pathologic side of normal tissue repair that results when the normal wound healing programs go awry. Successful resolution of tissue injury requires several highly coordinated pathways, and this research focuses on the interplay between these overlapping pathways: immune effectors, inflammatory mediators and fibroproliferation in the resolution of fibrosis. Previously we have successfully prevented, mitigated, and even reversed established fibrosis using vaccinia vaccination immunotherapy in two models of murine lung fibrosis. The mechanism by which vaccinia reverses fibrosis is by vaccine induced lung specific Th1 skewed tissue resident memory (TRMs) in the lung. In this study, we isolated a population of vaccine induced TRMs - CD49a^+^ CD4^+^ T cells - that are both necessary and sufficient to reverse established pulmonary fibrosis. Using adoptive cellular therapy, we demonstrate that intratracheal administration of CD49a^+^ CD4^+^ TRMs into established fibrosis, reverses the fibrosis histologically, by promoting a decrease in collagen, and functionally, by improving lung function, without the need for vaccination. Furthermore, co-culture of *in vitro* derived CD49^+^ CD4^+^ human TRMs with human fibroblasts from individuals with idiopathic pulmonary fibrosis (IPF) results in the down regulation of IPF fibroblast collagen production. Lastly, we demonstrate in human IPF lung histologic samples that CD49a^+^ CD4^+^ TRMs, which can down regulate human IPF fibroblast function, fail to increase in the IPF lungs, thus potentially failing to promote resolution. Thus, we define a novel unappreciated role for tissue resident memory T cells in regulating established lung fibrosis to promote resolution of fibrosis and re-establish lung homeostasis. We demonstrate that immunotherapy, in the form of adoptive transfer of CD49a^+^ CD4^+^ TRMs into the lungs of mice with established fibrosis, not only stops progression of the fibrosis but more importantly reverses the fibrosis. These studies provide the insight and preclinical rationale for a novel paradigm shifting approach of using cellular immunotherapy to treat lung fibrosis.

## Introduction

Pulmonary fibrosis is a devastating disease with no effective treatments to cure, stop or reverse the unremitting, fatal fibrosis. A critical barrier to treating this disease is the lack of understanding of the pathways leading to fibrosis as well as those regulating the resolution of fibrosis. Pulmonary fibrosis is the common end point for a diverse group of disorders such as chronic hypersensitivity pneumonitis, idiopathic pulmonary fibrosis, silicosis, radiation pneumonitis, and collagen vascular diseases^1,2^

Fibrosis is the pathologic side of normal tissue repair that results when the normal wound healing programs go awry. Successful resolution of tissue injury requires not only the activation of effector cells and the marked increase in synthesis and deposition of extracellular matrix (ECM), but also the deactivation of these effector cells and the clearance of excess ECM to allow return to normal lung structure and function^1^. Thus, fibrosis may result from deviations in one or several of these highly coordinated pathways and this research focuses on the interplay between these overlapping pathways: immune effectors, inflammatory mediators and fibroproliferation in the resolution of fibrosis.

Supporting the notion of local immune cells interacting with local native lung cells is our novel published data demonstrating the critical role of tissue resident memory (TRM) cells in the resolution of established lung fibrosis^3,4^. We have shown that fibrosis is due to a dysregulated immune response that directs unremitting fibroproliferation and that this pathogenic process can be prevented, arrested and even reversed by the establishment of a robust tissue memory T cell response in the lungs^3,4^. That is, the induction of robust Th1 TRM cells in the lungs can negatively regulate fibrosis and re-establish tissue homeostasis^3,4^.

While clearly there is a “point of no return” with regard to organ fibrosis, it has become equally clear that lung damage characterized by increased ECM, fibroproliferation, and chronic inflammation may not only be arrested but also may be reversed by allowing the normal pathways of wound repair *to do their job*^4,5^ . Upon lung infection, TRMs play a critical role in not only directing the anti-pathogen response but also in the resolution of inflammation leading to preservation of lung architecture and function^6–8^. In addition to playing a critical role in the recall response to pathogens, numerous studies have demonstrated the ability of TRMs to mediate protection against tissue specific challenges such as viral, bacterial, and parasitic infections ^6–8^. This protection is achieved by the interaction of TRMs with cells of the adaptive and innate immune response to collectively coordinate and promote immunity and to dictate the local inflammatory response^9,10^. This local response in turn may affect the recruitment, proliferation, and activation of not only immune cells but also resident lung cells.

The rationale behind the use of immunotherapy to treat lung fibrosis is based on the expanding literature concerning the role of inflammation in coordinating not only the immune response but also the response of resident cells in fibrosis^1^. Although it is the unregulated fibroproliferation that leads to fibrosis, the mechanisms of fibroblast/myofibroblast recruitment and activation remain poorly understood. Our group and others have identified a role for IL-17 and Th17 cells in promoting the inflammation leading to fibrosis^3,4,11–13^. In contrast, Th1 immune responses in the lung are associated with resolution of inflammation^3,4^. Thus, we sought a mechanism by which we could skew the lung environment away from the pro-fibrotic Th17 toward a pro-resolution Th1 environment. Given the successful use of immunotherapy in treating cancer, we sought to apply this knowledge to the lungs^3,14–18^. Indeed, we previously have successfully prevented, mitigated, and even reversed established fibrosis using immunotherapy in the form of vaccina vaccination in two models of murine lung fibrosis^3,4^. In this study, we further demonstrate that not only does vaccinia induce TRMs that can reverse fibrosis but so do other vaccines such as influenza vaccine (FluMist®). Furthermore, we have been able to isolate the population of vaccine induced TRMs – CD49a^+^ CD4^+^ T cells - that are both necessary and sufficient to reverse established pulmonary fibrosis. In fact, using adoptive cellular therapy we demonstrate that intratracheal administration of CD49a^+^ CD4^+^ TRMs into established fibrosis, reverses the fibrosis histologically, by promoting a decrease in collagen levels, and functionally, by improving lung function, without the need for vaccination. Furthermore, co-culture of *in vitro* derived CD49^+^ CD4^+^ human TRMs with human IPF fibroblasts results in the down regulation of idiopathic pulmonary fibrosis (IPF) fibroblast collagen production. Lastly, we demonstrate in human IPF lung histologic samples that CD49a^+^ CD4^+^ TRMs, which can down regulate human IPF fibroblast function, fail to increase in IPF lungs, thus potentially failing to promote resolution.

In this paper, we define a novel unappreciated role for tissue resident memory T cells in regulating established lung fibrosis to promote resolution of fibrosis and re-establish lung homeostasis. We demonstrate that immunotherapy, in the form of adoptive transfer of CD49a^+^ CD4^+^ TRMs into the lungs of mice with established fibrosis, not only stops progression of the fibrosis but more importantly reverses the fibrosis. These studies provide the insight and preclinical rationale for a novel paradigm shifting approach of using cellular immunotherapy to treat lung fibrosis.

## Materials and Methods

### Mice

*C57BL/6* were purchased from The Jackson Laboratory. 8 to 10-week-old female and male mice were utilized.

### Reagents

Modified vaccinia ankara virus was generated as previously described. This modified vaccinia ankara virus is a modified vaccinia that contains the full-length ovalbumin protein but lacks lytic ability. Quadrivalent FluMist® was purchased from AstraZeneca (Wilmington, DE). Bleomycin was purchased from App Pharmaceuticals (Schumburg, IL). PMA and Ionomycin were purchased from Sigma-Aldrich (St. Louis, MO). Flow cytometry reagents were purchased from BD Biosciences (Franklin Lakes, NJ). Antibodies utilized were from Biolegend (San Diego, CA), see Supplemental Table 1.

### Cell Lines

Age matched normal and IPF lung fibroblasts were purchased from BioIVT (Westbury, NY). Fibroblasts were maintained in DMEM supplemented with 10% FBS. For all coculture experiments fibroblasts were used between passages 3-6. To generate human CD49a^+^ and CD49a^-^ CD4^+^ T cells, lymphocytes were isolated from buffy coats of normal donor leukopaks. Bulk lymphocytes were stimulated with anti-CD3/anti-CD28 (Miltenyi Biotech) in RPMI media supplemented with 10% FBS, antibiotic, and glutamine. Two days later human IL2 (Peprotech, East Windsor, NJ), was added at 10 ng/mL. Cultures were maintained for 4 weeks with addition of media as needed and addition of IL2 every 7 days.

### Coculture Experiments

Fibroblasts were harvested and 50,000 were plated per well of a 24 well transwell plate (Corning, Corning, NY.) Fibroblasts were allowed to adhere for 6 hours before addition of T cells. Live human lymphocytes in culture were isolated by Ficoll-Paque (Fisher Scientific, Waltham, MA). CD4^+^ T cells were then isolated by negative magnetic isolation (Biolegend). CD49a^+^ and CD49a^-^ cells were isolated by magnetic isolation using Anti-CD49a purified antibody (Biolegend) and anti-mouse IgG microbeads (Miltenyi Biotech). Purified CD4 T cells were added into transwells of 24 well plates in DMEM media supplemented with 10pg/mL human IL2. Cocultures were incubated overnight, followed by addition of 10ng/mL TGFβ (Peprotech). 24 hours later fibroblast RNA was harvested.

### Flow cytometry

All Flow cytometry analysis was performed on a BD FACSCelesta (BD Biosciences) and analyzed using FlowJo software (TreeStar Inc, Ashland, OR).

### Pulmonary fibrosis model

Mice were injected i.p. with 0.8 units bleomycin on days 0, 3, 7, 14, 21, and 28 to induce pulmonary fibrosis. For some experiments mice received bleomycin intratracheally at a dose of .018 units in 25ul PBS.

### Vaccination

Vaccinia vaccine was administered intranasally at a dose of 2 million pfu per mouse. Quadrivalent FluMist® was administered intranasally at a dose of 10^5.5–6.5^ fluorescent focus units of live attenuated influenza virus reassortants of each of the four strains: A/Victoria/1/2020 (H1N1) (A/Victoria/2570/2019 (H1N1) pdm09-like virus), A/Norway/16606/2021 (H3N2) (A/Darwin/9/2021 (H3N2)-like virus), B/Phuket/3073/2013 (Yamagata lineage), and B/Austria/1359417/2021 (Victoria lineage).

### ELISA

TNFα and IFNγ ELISA were purchased from ebioscience (San Diego, CA), and performed according to manufacturer’s instructions.

### Histology

Lungs were inflated to 27cm H_2_0 with 10% neutral-buffered formalin, sectioned and stained for H&E and Masson’s trichrome according to previously established procedures^19^. Samples were analyzed by microscope at 40x magnification.

### RNA Extraction and Real Time PCR

Total lung RNA was extracted using TRIzol reagent (Invitrogen, Carlsbad, CA) and reversed transcribed by Reliance (Bio-Rad Laboratories, Hercules, CA) as per the manufacturer’s protocol. Real-time PCR was performed on an Applied Biosystems 7300 PCR machine using Applied Biosystems reagents (Carlsbad, CA) and normalized to 18s rRNA. Values were calculated using the delta Ct method in reference to control samples for each primer. All primers and probe sets used were purchased from Applied Biosystems (Carlsbad, CA).

### RNAseq analysis

Wild type mice were vaccinated intranasally with FluMist, and 4 weeks later *in vivo* labeled CD49a^+^ CD4^+^ lung TRM and CD49a^-^ CD4^+^ lung TRM were sorted into Trizol. RNA was isolated and subjected to RNA seq analysis. Alignments were performed using Hisat2, and differential gene expression was performed using DESeq2 and BiomaRt. The top 500 most differentially expressed genes in CD49a^+^ CD4^+^ lung TRMs were submitted to STRING analysis of known and predicted protein-protein interactions. Clustering of upregulated proteins resulting in a cluster which contained 25 upregulated genes with a small PPI enrichment p-value (< 10e-16), indicating a high likelihood that the proteins are biologically connected.

### Analysis of scRNAseq data

Processed scRNAseq data for human lungs with fibrosis were obtained from the Gene Expression Omnibus database (accession number GSE16831)^20^. Annotated T cells were clustered using the R software package Seurat (v.4.2). CD49a^+^ TRM were identified by CD4, CD69, and ITGA1 (CD49a) expression (log-transformed expression levels > 0.5). CD103a^+^ TRM were identified by CD4, CD69, and ITGAE (CD103) expression. TRMs were plotted as % of annotated T cells in IPF (n=17) and non-fibrotic control (n=10) lungs. Lungs with fewer than 200 CD4^+^ T cells or with no CD49a^+^ or CD103^+^ CD4+ T cells were excluded.

### In vivo antibody labeling and sorting

For *in vivo* antibody labeling, mice were injected i.v. with 2.5 μg PE-conjugated anti-CD4 antibody (clone RM4-5), and after 10 minutes, lungs were isolated, rinsed in PBS, and digested in a mixture of 3mg/mL Collagenase and Dnase I for 45 minutes at 37°C. Cells were filtered into single cell suspensions, and CD4^+^ T cells were positively selected by magnetic isolation (Miltenyi Biotech (Auburn, CA). Isolated lymphocytes were then stained *in vitro* with a different, noncompeting clone of APC–anti-CD4 (clone RM4-4), along with antibodies to other surface markers with fluorochrome-conjugated antibodies. Stained cells were sorted using a BD FacsAria (BD Biosciences, Franklin Lakes, NJ). CD4^+^ T cells which were singly stained with APC anti-CD4 *in vitro* but not stained with PE anti-CD4 *in vivo* were considered to be tissue resident. For TRM transfer experiments, sorted T cells were resuspended in PBS at 1x10^6^ per ml and mice received 50ul intratracheally.

### Diffusion factor for CO measurement

To assess the efficiency of gas exchange in the lungs following bleomycin-induced injury, measurement of the diffusion factor for CO (DFCO) was performed as described previously^21^. Briefly, mice were anesthetized with a mixture of ketamine (100 mg/kg)/xylazine (15 mg/kg) via i.p. injection. Once sedated, mice were intubated with a 20-gauge IV angiocatheter. Mouse lungs were quickly inflated with a 0.8 ml gas mixture (0.5% neon, 1% CO and 98.5% air). After a 9-second breath hold, 0.8 m of gas was quickly withdrawn from the lung and diluted to 2 ml with room air. The neon and CO concentrations in the diluted air were measured by gas chromatography (INFICON, Model 3000A) to assess DFCO. The dilution to 2 ml was needed, since the gas chromatograph required a minimal sample size of 1 ml.

### Pulmonary Mechanics Measurements

After DFCO assessment, mice were connected to a flexi-Vent ventilator (SCIREQ) and ventilated with a tidal volume of 0.2 ml of 100% oxygen at a rate of 150 Hz. with a positive end–expiratory pressure (PEEP) of 3 cmH_2_O. Mice were subjected to deep inspiration at 30 cmH_2_O for 5 seconds and returned to normal ventilation for 1 minute. Baseline measurements of respiratory system resistance (Rrs), compliance (Crs) and elastance (Ers) were measured using forced oscillation technique via the SnapShot perturbation, a single 2.5-Hz sinusoidal waveform which is fit to the single compartment model via linear regression. Following measurements angiocatheters were removed and mice were observed until they woke up from anesthesia.

### Immunofluorescence Staining

Formalin-fixed, paraffin-embedded (FFPE) human IPF and normal tissue slides were incubated with primary antibody to CD4, CD49a, and CD103 in 5% goat serum, 0.5% BSA, in PBS at 4°C overnight. Images were collected at x20 magnification using a Nikon Eclipse 80i microscope and Nikon DS-fi1 camera.

### Statistical Analysis

All experiments were performed in biological triplicate and results represent the mean ± standard deviation. All experiments were replicated at least three times. Statistical analysis was performed using either 1-way ANOVA followed by Tukey’s test or a paired Student’s *t*-test; *p*<0.05 was considered statistically significant.

## Results

### CD49a and CD103 expression differentiates populations of lung CD4^+^ TRM

We previously demonstrated that immunotherapy in the form of intranasal vaccinia virus induced a pool of lung, tissue resident, memory Th1 CD4^+^ T cells in mice^3,4^. We also demonstrated that generation of lung TRM was required to mitigate bleomycin induced pulmonary fibrosis^3,4^. To further understand the effect of immunotherapy induced lung TRM in mice we intranasally administered PBS, quadrivalent FluMist®, or vaccinia virus. To analyze TRM generation during the development of fibrosis we also treated a group of mice with intratracheal bleomycin. Twenty-eight days later we intravenously labeled circulating T cells with CD4 antibody to identify and isolate lung tissue resident CD4^+^ T cells. Mice were sacrificed and lungs were removed, processed to single cell suspensions, and stained for CD4 and the surface markers CD49a and CD103. Lung CD4^+^ T cells which were not stained *in vivo* and thus only singly stained with CD4 *in vitro* were considered tissue resident. CD49a^+^ and CD103^+^ CD4^+^ lung TRM were sorted and stimulated for 4 hours to perform intracellular cytokine staining (ICS).

TRM from bleomycin treated mice were predominantly CD103^+^, whereas TRM from vaccinated mice were predominantly CD49a^+^ (Figure 1a, b). PBS treated mice also had predominantly CD49a^+^ TRM, however the percentages and total numbers of these cells were significantly lower than vaccinated mice. ICS analysis of sorted CD4^+^ TRM demonstrated increased expression of IL17a and minimal expression of IFNγ by CD103^+^ TRM regardless of how mice were treated (Figure 1c). ICS analysis of CD49a^+^ TRM demonstrated decreased IL17a production relative to CD103^+^ TRM regardless of treatment, but only increased IFNγ production in mice that received either vaccinia or FluMist® vaccination. Our previous data indicated that IFNγ production was required for the therapeutic effect observed by vaccination induced TRM^3,4^. Based on this finding and these ICS data, we hypothesize that CD49a^+^ CD4^+^ TRM are responsible for the therapeutic effect of vaccination on fibrosis resolution in the lung.

**Figure 1.**
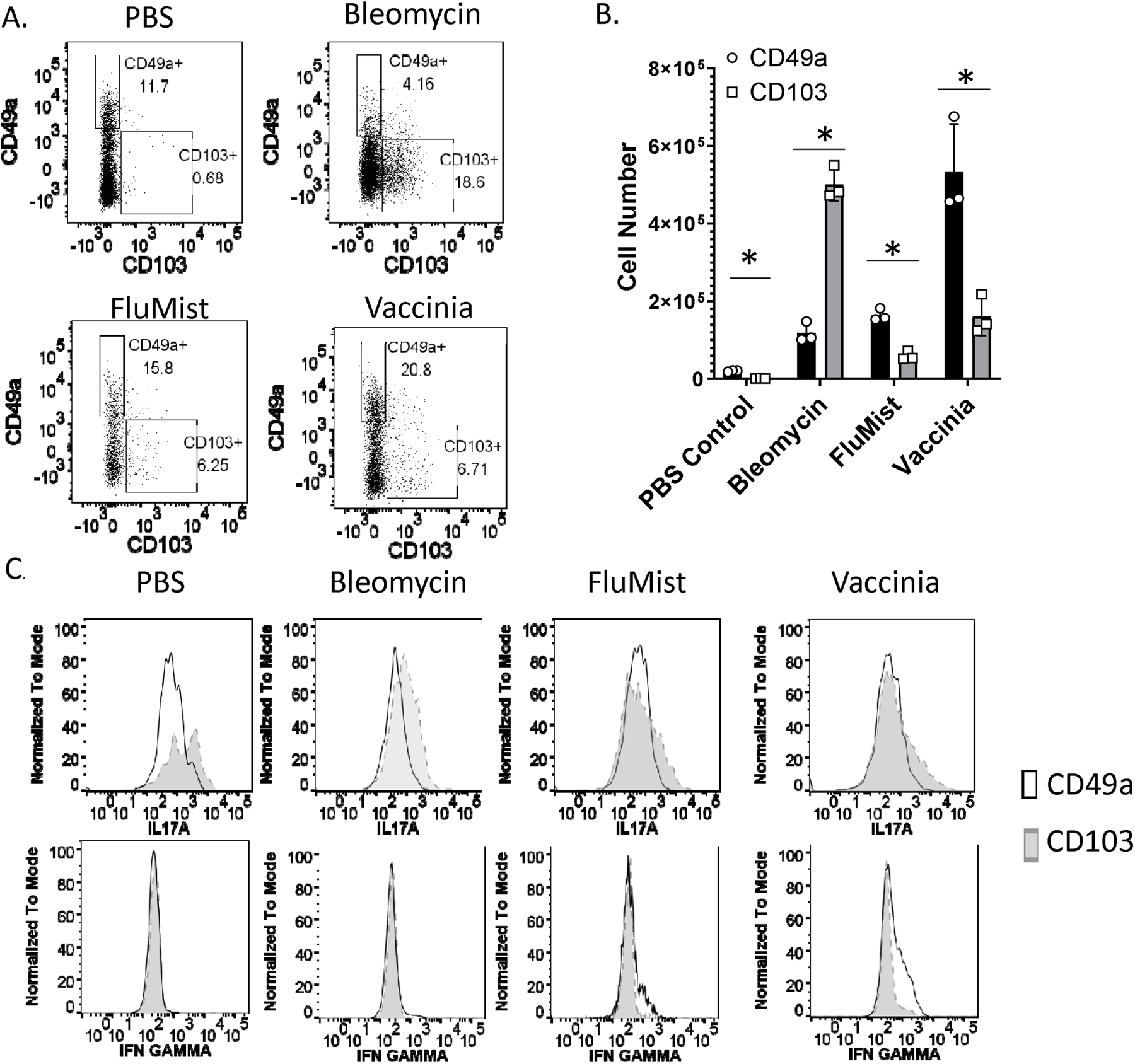
CD49a and CD103 expression differentiates populations of lung CD4^+^ TRM. Mice received intranasal PBS, intratracheal bleomycin, intranasal FluMist®, or intranasal vaccinia virus. 28 days later mice were injected i.v. with CD4 antibody in order to identify tissue resident CD4^+^ T cells, sacrificed, and lungs were isolated. **A.** Flow plots of tissue resident CD4^+^ T cells stained with CD49a and CD103 antibodies **B**. Numbers of lung CD49a^+^and CD103^+^ CD4^+^ TRM. **C.** Intracellular staining of IL17a and IFNγ from sorted CD49a^+^ and CD103^+^ lung CD4^+^ TRM. Error bars represent standard deviation. * indicates statistical significance where p < .05. Experiments were performed three times with 3 mice per group.

### CD49a**^+^** CD4**^+^** TRM are therapeutic in pulmonary fibrosis

To further analyze the effect of CD49a^+^ CD4^+^ TRM on fibrosis we injected mice intraperitoneally with bleomycin on days 0, 3, 7, 14, 21, and 28 to induce pulmonary fibrosis. Unlike the intratracheal model of pulmonary fibrosis, this model results in long term progressive lung fibrosis which better mimics human disease ^5^. To generate TRM we intranasally vaccinated another cohort of mice with FluMist® and 28 days later sorted lung CD4^+^ TRM into two populations, CD49a^+^ CD4^+^ T cells, and CD49a^-^ CD4^+^ T cells (figure 2a). We intratracheally transferred 50,000 sorted CD49a^+^ or CD49^-^ CD4^+^ T cells into the IP bleomycin treated mice 42 days after the initiation of bleomycin treatment. Thirty days later, (day 72), mice underwent pulmonary function testing. Mice that received CD49a^-^ CD4^+^ TRM demonstrated increased tissue resistance, decreased compliance, and decreased diffusion capacity of carbon monoxide (CO) relative to PBS treated control mice (figure 2b). In contrast, mice which received CD49a^+^ CD4^+^ TRM demonstrated no notable change in tissue resistance, lung compliance or diffusion capacity relative to PBS control mice, indicating a therapeutic role for these TRM in preventing the progression of fibrosis but more importantly in promoting the resolution of the fibrosis. When we analyzed lung RNA for collagen expression, we observed significantly increased levels of collagen 1 and collagen 3 in the lungs of mice which received CD49a^-^ CD4^+^ TRM. However, lungs from mice which received CD49a^+^ CD4^+^ TRM failed to demonstrate a significant increase in collagen 1 or collagen 3 relative to PBS treated mice. These data again support a therapeutic role for CD49a^+^ CD4^+^ TRM in preventing the progression of pulmonary fibrosis (Figure 2c).

**Figure 2.**
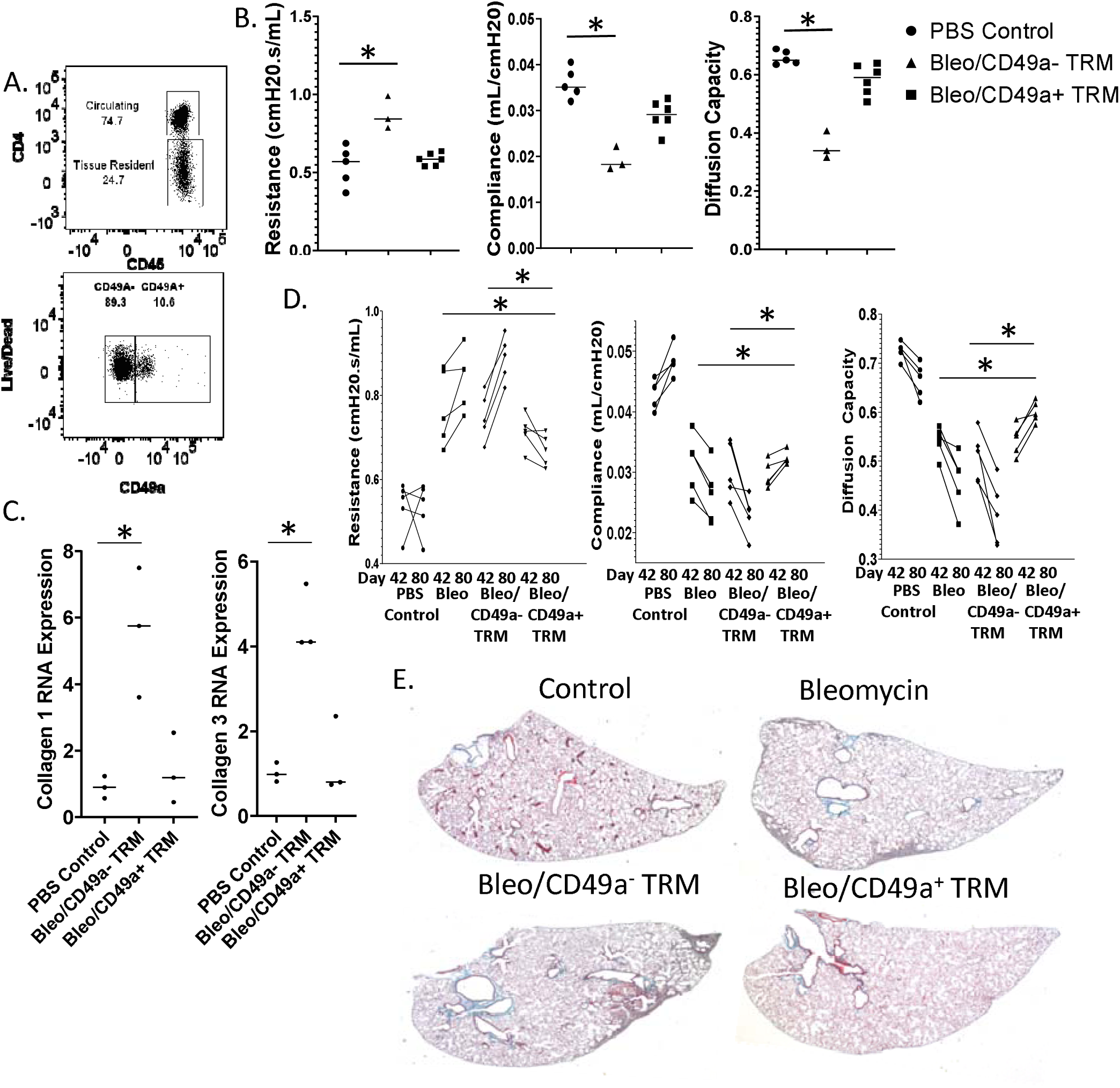
CD49a ^+^ CD4^+^ TRM are therapeutic in pulmonary fibrosis. Pulmonary fibrosis was established in mice using intraperitoneal bleomycin injections. CD49a^+^ CD4^+^ T cells, and CD49a^-^ CD4^+^ T cells were sorted from lungs of mice which had received intranasal FluMist® 28 days prior. Sorted TRM were transferred intratracheally into mice with established pulmonary fibrosis on day 42 of the I.P. bleomycin model. Thirty days later mice were subjected to pulmonary function testing. **A.** Flow plots of sorted CD4^+^ TRM prior to transfer. **B.** Pulmonary function testing of I.P. bleomycin treated mice. **C.** Lung Collagen RNA expression 45 days following T cell transfer. **D.** Changes in pulmonary function from day 42 (pre-TRM transfer) and day 80 (post-TRM transfer). **E.** Represent histology of I.P. bleomycin treated mice sacrificed on day 100. Error bars represent standard deviation, * indicates statistical significance where p < .05. Experiments were performed three times with 8 mice per group.

To further confirm the therapeutic role of CD49a^+^ CD4^+^ T cells in our model of bleomycin and to rule out unforeseen effects on mice lung health or experimental variability from bleomycin injections, we performed a time course experiment. We again induced pulmonary fibrosis by repeat I.P. injections of bleomycin and then performed baseline pulmonary function testing on mice at day 42 before any adoptive transfer of TRMs. Only mice which demonstrated decreased pulmonary function as determined by increased tissue resistance, decreased lung compliance, and decreased diffusion capacity were selected for CD4^+^ T cell transfer. To induce CD4^+^ T cells for transfer, a different cohort of mice received intranasal FluMist® and 4 weeks later lung CD49a^+^ CD4^+^ TRM and CD49a^-^ CD4+ TRM were isolated by sorting. On day 49, mice with decreased lung function received either no CD4^+^ T cells, 50,000 CD49a^-^ CD4^+^ TRM, or 50,000 CD49a^+^ CD4^+^ TRM intratracheally. Mice were followed for 31 more days and on Day 80, pulmonary function testing was again performed. Mice that received bleomycin without T cell transfer or bleomycin followed by CD49a^-^ CD4^+^ TRM demonstrated increased resistance, decreased lung compliance, and decreased diffusion capacity relative to their function observed at day 42 and to PBS controls, consistent with the continued development of lung fibrosis (figure 2d). However, mice which received bleomycin followed by CD49a^+^ CD4^+^ TRM demonstrated decreased tissue resistance, increased lung compliance and increased diffusion capacity at day 80 relative to their function observed at day 42. More specifically, the mice that received CD49a^+^ TRM had improved lung function between days 42 and 80 indicating reversal of the progressive fibrosis seen in the other experimental conditions (Figure 2d). Histology of lungs isolated at day 100 confirmed our pulmonary function data, with lungs from mice which received no T cells or CD49a^-^ CD4^+^ TRM demonstrating increased collagen deposition and cellular infiltration, whereas lungs from mice which received CD49a^+^ CD4^+^ TRM resembled lungs from PBS treated mice (figure 2e). These data indicate that even after the initiation of lung fibrosis as determined by increased resistance, decreased compliance, and decreased diffusion capacity observed at day 42, transfer of CD49a^+^ CD4^+^ TRM is capable of halting and reversing the development of pulmonary fibrosis as demonstrated by the improvements of lung function observed at day 80.

### Human CD49a**^+^** CD4**^+^** T cells suppress profibrotic gene expression

Having shown that CD49a^+^ CD4^+^ T cells were therapeutic in our bleomycin model of pulmonary fibrosis, we wanted to determine if this was also occurring in human idiopathic pulmonary fibrosis. To accomplish this, we elected to perform coculture experiments with human CD4^+^ T cells and lung fibroblasts cultured from biopsies of IPF patients. We felt that this experimental design was appropriate since we had observed decreased levels of collagen 1 and 3 expression in mice which received CD49a^+^ CD4^+^ T cells, and lung fibroblasts are known to be significant producers of collagen in the fibrotic lung.

To generate CD49a^+^ CD4^+^ T cells, human PBMCs were stimulated with anti-CD3 and anti-CD28 in the presence of interlukin-2 for four weeks. CD4^+^ T cells were magnetically isolated and then the cells were magnetically separated based on CD49a expression. Stimulation of PBMCs resulted in the upregulation of CD49a on approximately 25% of isolated CD4^+^ T cells, and CD49a^+^ T cells were effectively isolated by magnetic separation (Figure 3a). To characterize human CD49a^+^ CD4^+^ T cells, we stimulated isolated CD4^+^ T cells with PMA and Ionomycin and performed ICS analysis. Functionally CD49a^+^ CD4^+^ T cells produced more of the classical Th1 cytokines TNFα, granzyme b, and IFNγ than CD49a^-^ CD4^+^ T cells(Figure 3b).

**Figure 3.**
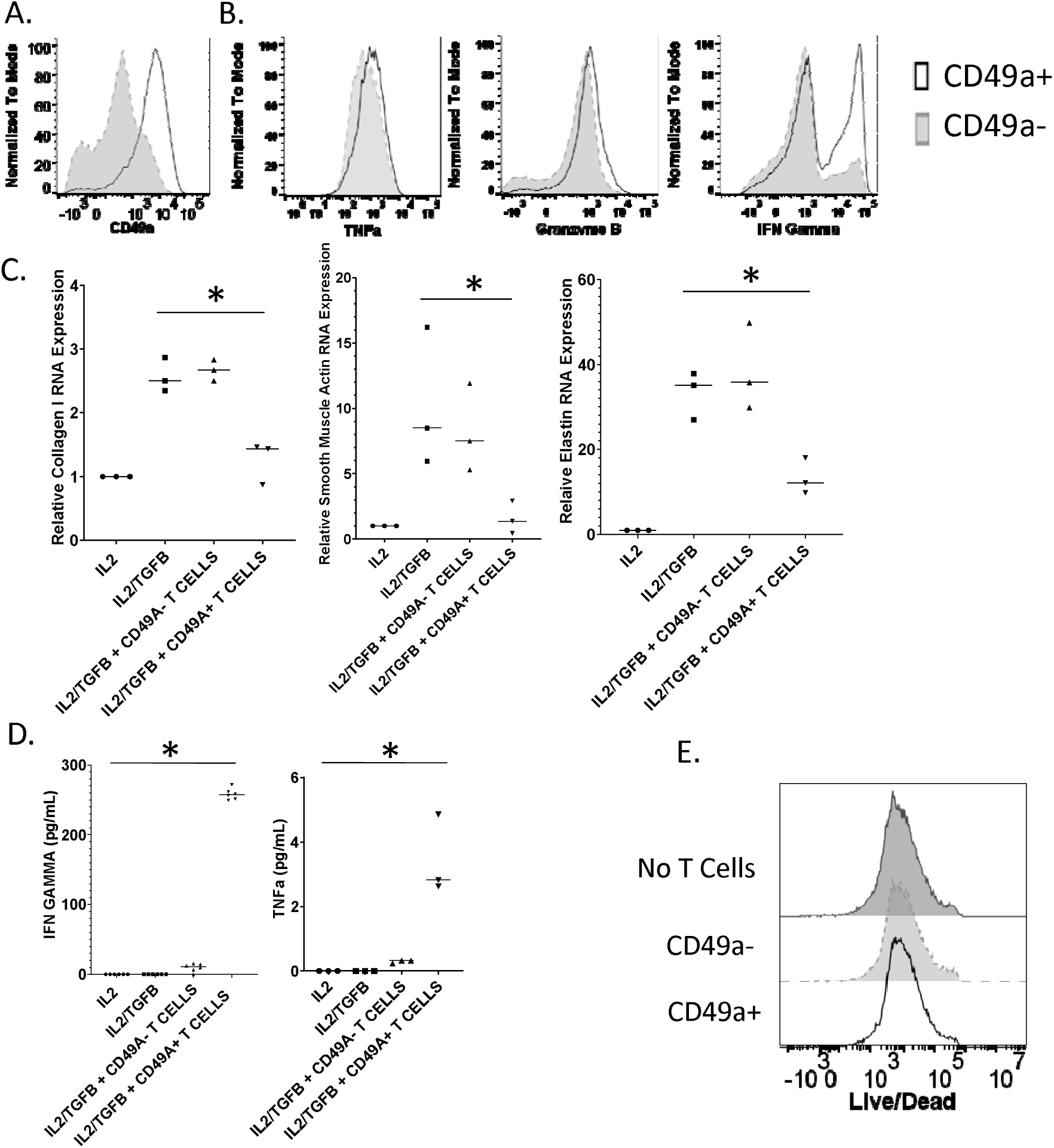
Human CD49a^+^ CD4 ^+^ T cells suppress lung fibroblast profibrotic gene expression. In order to generate CD49a^+^ CD4^+^ T cells, human PBMCs were stimulated with anti-CD3 and anti-CD28 in the presence of interlukin-2 for four weeks. CD4^+^ T cells were isolated, and cells were separated based on CD49a expression. **A.** Flow analysis of CD49a expression on isolated human CD4^+^ T cells. **B.** Flow analysis of cytokine production by CD49a^-^ or CD49a^+^ CD4^+^ T cells. **C.** RNA expression of pro-fibrotic genes by IPF lung fibroblasts cocultured with CD49a^-^ or CD49a^+^ CD4^+^T cells in transwell plates. **D.** TNFα and IFNγ concentration of coculture supernatants measured by ELISA. **E.** Live/Dead staining of IPF fibroblasts following no coculture, or coculture with CD49a^-^ and CD49a^+^ CD4^+^ T cells. Error bars represent standard deviation, * indicates statistical significance where p < .05. Experiments were performed three times.

To assess the effect of the isolated CD4^+^ T cells on IPF fibroblasts, transwell coculture experiments were performed where CD4^+^ T cells were added at a 2:1 ratio with lung fibroblasts from IPF patients. IL2 was added to stimulate the CD4^+^ T cells and cells were incubated overnight before the addition of TGFβ to induce profibrotic gene expression. Fibroblast RNA was collected after 24 hours of TGFβ stimulation. The addition of TGFβ induced a significant increase in collagen 1, smooth muscle actin, and elastin gene expression (Figure 3c). Coculture with CD49a^-^ CD4^+^ T cells had no effect on profibrotic gene expression relative to TGFβ addition alone, however coculture with CD49a^+^ CD4^+^ T cells resulted in a significant reduction in profibrotic gene expression supporting a role for these T cells in the setting of decreasing fibrosis. When we assayed coculture supernatants for cytokines we observed increased levels of TNFα and IFNγ in wells to which CD49a^+^ CD4^+^ T cells were added. While overall TNFα levels were barely detectable, the levels of IFNγ were significantly increased in wells which contained CD49a^+^ CD4^+^ T cells, confirming our ICS analysis data (Figure 3d). Since we observed increased granzyme b levels following stimulation of CD49a^+^ CD4^+^ T cells we elected to determine if the observed decrease in profibrotic gene expression was due to killing of IPF fibroblasts. To accomplish this IPF fibroblasts were isolated post coculture and stained with a live/dead dye and flow cytometry was performed. Coculture with either CD49a^-^ or CD49a^+^ CD4^+^ T cells did not result in increased death of IPF fibroblasts as determined by equivalent exclusion of dye (Figure 3e). These data support a therapeutic role for CD49a^+^ CD4^+^ T cells against fibrotic progression and specifically demonstrate the ability of human CD49a^+^ CD4^+^ T cells to suppress profibrotic gene expression by IPF fibroblasts.

### CD49a+ TRM alter the fibrotic lung environment

Although we have demonstrated that in mice intratracheally transferred CD49a^+^ CD4^+^ TRM can improve lung function, reverse established fibrosis *in vivo,* and can suppress human fibroblast profibrotic gene expression *in vitro*, we wanted to determine how these TRMs were modifying the fibrotic lung environment. To accomplish this, we vaccinated wild type mice with intranasal FluMist® and 4 weeks later sorted CD49a^+^ CD4^+^ TRM and CD49a^-^ CD4^+^ TRM and then performed RNA sequence analysis. Differential gene analysis comparing lung CD49a^+^ CD4^+^ TRM to lung CD49a^-^ CD4^+^ TRM demonstrated significantly increased expression of the extracellular matrix modifying genes Adamts1, 2, 4, 5, 8, 9, 12, 15, and 17. In addition, we observed decreased expression of the cytokines IL17a and IL17f consistent with the phenotype we observed previously (Figure 4a). To identify upregulated molecular pathways, we submitted the top 500 most significantly upregulated differentially expressed genes by CD49a^+^ CD4^+^ T cells to STRING analysis of known and predicted protein-protein interactions. Clustering of upregulated proteins resulting in a cluster which contained 25 upregulated genes with a small PPI enrichment p-value (< 10e-16), indicating a high likelihood that the proteins are biologically connected (Figure 4b). This cluster contained many proteins involved in extracellular matrix and collagen binding. In addition, the cluster contained several proteases and metallopeptidases. These data suggest that CD49a^+^ CD4^+^ TRM are acting directly on the fibrotic extracellular matrix, potentially binding to deposited collagen. Through this binding, CD49a^+^ CD4^+^ T cells may be inducing degradation of collagen and other extracellular matrix proteins present in increased quantities during fibrosis through the expression of metallopeptidases.

**Figure 4.**
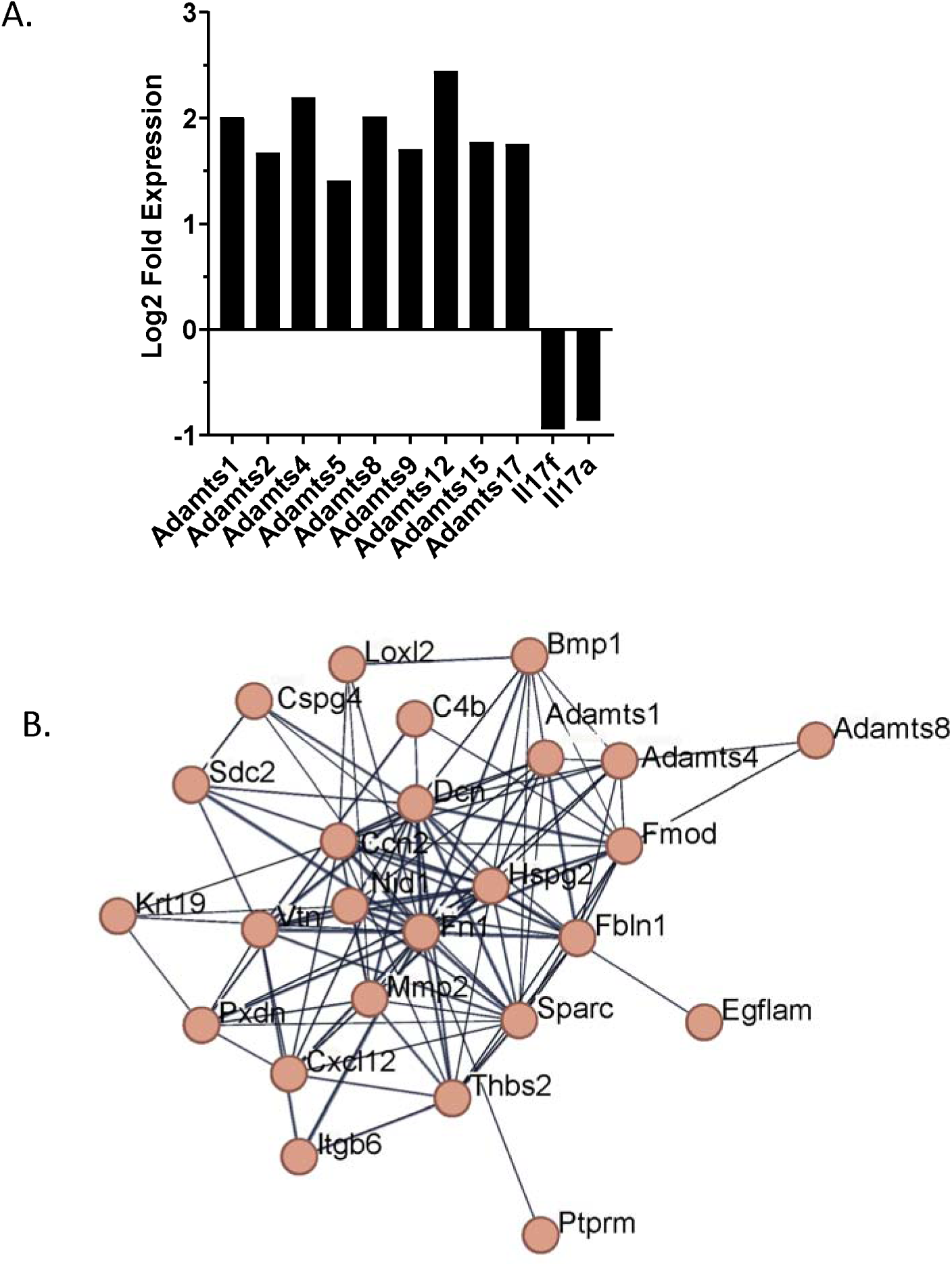
CD49a ^+^ CD4 ^+^ TRM gene signature indicates the ability to bind and alter the extracellular matrix. CD49a^-^ and CD49a^+^ CD4^+^ T cells were sorted from lungs of mice which had received intranasal FluMist® 28 days prior. RNA was isolated and subjected to RNA seq analysis. **A.** STRING clustering figure of the 500 most differentially expressed genes by CD49a^+^ CD4^+^ TRM. **B.** Differential gene expression analysis of CD49a^+^ CD4^+^ TRM versus CD49a^-^ CD4^+^ TRM.

### Presence of CD49a**^+^** CD4**^+^** T cells in healthy and fibrotic human lungs

We have observed that human CD49a^+^ CD4^+^ T cells are capable of suppressing profibrotic gene expression *in vitro* (Figure 3), and that transfer of murine CD49a^+^ CD4^+^ T cells can both prevent further lung fibrosis as well as reverse established fibrosis in bleomycin treated mice (Figure 2). We have also observed that bleomycin treated mice have an increase in CD103^+^ CD4^+^ TRM relative to CD49a^+^ CD4^+^ TRM, whereas control and vaccinated mice have ratios favoring CD49a^+^ CD4^+^ TRM to CD103^+^ CD4^+^ TRM (Figure 1b). These findings indicate that during fibrosis there is a skewing of lung TRM towards CD103^+^ expression and away from CD49a^+^ expression, which we believe represents a pathologic phenotype. As such, our ability to reverse this skewing by adoptive transfer of CD49a^+^ CD4^+^ TRM may be a therapeutic phenotype. As such, we were interested in determining if this skewing of lung TRM cells was also present in human lungs. For this, we analyzed scRNAseq data from human lungs obtained from the Gene Expression Omnibus database (accession number GSE16831) ^20^. Analysis of the data from annotated T cells indicated that there were significantly more CD103^+^ TRMs in IPF lungs compared to control lungs, whereas the percentage of CD49a^+^ TRMs was relatively unchanged (Figure 5a). Thus, (at least based on transcriptomic data), it appears that there is a similar skewing of human lung CD4^+^ TRMs favoring an increase in CD103^+^ CD4^+^ TRMs relative to CD49a^+^ CD4^+^ TRMs in human IPF. To further assess the presence of these cells in healthy and diseased lungs, we performed immunofluorescence on lung biopsy samples from age matched normal controls and IPF patients. IHC of CD4 and CD103 demonstrated few CD103^+^ CD4^+^ T cells in normal lungs (Figure 5b). However, IHC of IPF lungs showed large numbers of CD4^+^ T cells, many of which co-stained with CD103. In contrast, IHC of CD4 and CD49a demonstrated populations of CD49a^+^ CD4^+^ T cells in normal lungs, but few CD4^+^ T cells co-staining with CD49a in the lungs of IPF patients. These data strongly support a similar increase in CD103^+^ CD4^+^ TRMs relative to CD49a^+^ CD4^+^ TRMs in human IPF, mirroring the fibrotic animal phenotype that was reversed with cellular immunotherapy (Figure 2). Overall, these data indicate that there is an alteration in the phenotype of lung TRMs during fibrosis which permits the progression of this disease state and that transfer of CD49a^+^CD4^+^ TRM into the lung represents a therapeutic intervention to swing the balance back toward homeostasis.

**Figure 5.**
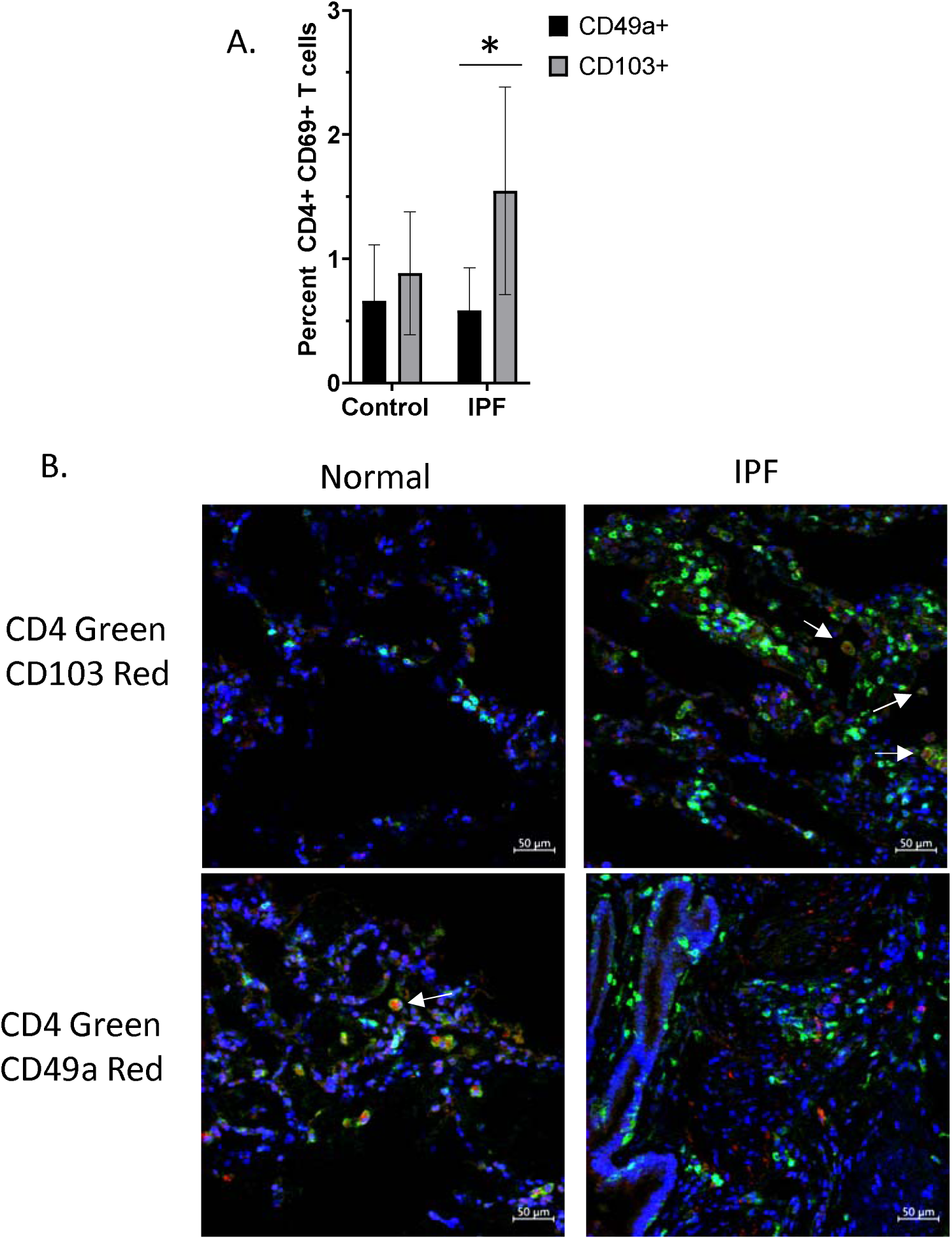
Imbalanced CD49a^+^/CD103^+^ CD4^+^ T cells in human IPF lungs. **A** Percentage of CD49a^+^ and CD103^+^ CD4^+^ T cells in control and IPF lungs as determined by analysis of single cell RNA-seq dataset from Adams et al (2020). Error bars represent standard deviation, * = p value of 0.0001. **B.** Immunohistochemistry was performed on biopsies of normal and IPF human lungs in order to determine densities of CD49a^+^ and CD103^+^ CD4^+^ T cells. Images were taken at 40X.

## Discussion

In this paper, we demonstrate that immunotherapy, in the form of adoptive transfer of CD49a^+^ CD4^+^ TRMs into the lungs of mice with established fibrosis, not only stops progression of fibrosis but more importantly reverses fibrosis. We define a novel unappreciated role for tissue resident memory T cells in regulating pathologic fibrosis to promote resolution of fibrosis and re-establish lung homeostasis. These studies provide the insight and preclinical rationale for a novel paradigm shifting approach of utilizing immunotherapy for lung fibrosis in general and specifically IPF, a deadly disease with no treatment to stop or reverse fibrosis.

Current medications for IPF at best slow down the fibrosis without impacting mortality or quality of life^22–24^. Although the exact cause of the disease is unknown, it appears to be related to dysregulated wound healing. The immune microenvironment in the lung is constantly exposed to the external environment and thus has developed a robust immune response to potential threats. As such, alterations in this response may result in excessive responses and lack of appropriate downregulation and healing^25^.

We have previously demonstrated in a chronic mouse model of lung fibrosis that immunotherapy, in the form of vaccinia vaccination, not only prevents and halts lung fibrosis, but it also actually reverses established lung fibrosis^3,4^. The mechanism by which vaccinia reverses fibrosis is via vaccine induced lung specific Th1 skewed TRMs in the lung^3,4^. In this study, we further demonstrate that not only does vaccinia induce TRMs that can reverse fibrosis but so do other vaccines such as an influenza vaccine (FluMist®). Furthermore, we now have been able to isolate the exact vaccine induced TRMs that are both sufficient and necessary, CD49a^+^ CD4^+^, to reverse the established fibrosis. In fact, using adoptive cellular therapy we demonstrate that intratracheal administration of CD49a^+^ CD4^+^ T cells into established fibrosis, reverses the fibrosis histologically, by promoting a decrease in collagen levels, and functionally without the need for vaccination. In addition, we demonstrate the paucity of these cells in histologic samples from patients with IPF as well as that co-culture of *in vitro* derived CD49a^+^ CD4^+^ TRMs with human IPF fibroblasts results in a down regulation of IPF fibroblast collagen production.

No mouse model of fibrosis is perfect, however in our model, chronic fibrosis is induced with IP injections of bleomycin over four weeks leading to progressive low-grade immune cell infiltration, collagen deposition and fibrotic changes in the mouse lungs at 72^+^ days^3,4^. This model differs from an acute IT bleomycin model because it never causes an acute inflammatory phase but rather promotes sub-acute inflammation that progresses to fibrosis over months^3,4,26,27^. As an acute inflammatory stage is often unrecognized in humans and the lung fibrosis progresses slowly and continuously over time, the IP bleomycin model appears to more closely approximate human disease. Although numerous animal studies using the <u>acute</u> <u>intratracheal</u> bleomycin model of lung fibrosis have demonstrated benefit from several therapeutic agents, these often have not translated into human disease^28,29^. This may be in part be due to the fact that the intratracheal bleomycin model causes <u>acute</u> lung injury that heals with variable fibrosis^28–30^. This model is often neither progressive beyond the first few weeks and has been demonstrated to be reversible without any intervention over time^28,29^. Indeed, most drug interventions in this model are tested during this acute/subacute injury and may indeed only reflect accelerated resolution of lung injury thus preventing fibrosis. In humans, IPF likely develops over many years prior to presentation. Simply put, human disease is not temporally associated with acute lung injury and is also not punctuated with recurrent acute episodes in the majority of patients^31^. Our model of chronic lung fibrosis better approximates the relentless, progressive, human disease. In our model, fibrosis arises after subacute insults of intraperitoneal bleomycin over a month, followed by progressive fibrosis (in the absence of continued bleomycin insult) over the subsequent months ^3^. When we therapeutically intervene, it is temporally far removed from any acute phase. In fact, we quantitate the levels of lung fibrosis via survival pulmonary function testing on all mice on <u>day 42</u> after bleomycin and then randomize the mice to therapy with adoptive transferred CD49a^+^ TRM or CD49a^-^ TRM. Follow up is on <u>day 80</u> with repeat lung function testing, histology, FACS and cytokine expression. Thus, our data show that immunotherapy via adoptive transfer of CD49a^+^ TRM reverses established lung fibrosis in mice.

Although some memory T cells circulate in the blood and amongst secondary lymphoid organs as effector memory cells (Tem), memory T cells also take up permanent residence in specific tissue compartments^32–35^. These tissue resident memory cells (TRM), generated in response to site specific infections in lungs, skin, etc., are non-migratory and specifically maintained in these tissues^32–35^. Numerous studies have demonstrated an essential role for TRM in the recall response to mediate protection both against tissue specific and non-specific challenges such as viral, bacterial, and parasitic infections^6–8^. For example, in the lungs both CD4^+^ and CD8^+^ TRM are important for protection against influenza^8,9^. Interestingly, although both injectable inactivated influenza vaccine (IIV) and intranasal live attenuated influenza vaccine (LAIV) generate neutralizing strain-specific antibodies, only the intranasally administered LAIV generates lung localized, virus specific T cell responses similar to what is generated with influenza infection^8^. More importantly, only the intranasal LAIV generates TRM that mediate cross-strain protection, independent of circulating T cells and neutralizing antibodies^8^. Thus, intranasal LAIV generation of lung TRM not only protects from future infection to the specific vaccinated viral strain but also provides heterosubtypic protection to non-vaccine viral strains^8^.

In this regard, we too have published that only intranasal vaccination of live vaccinia generates TRMs that are essential both to the protection from and the reversal of bleomycin-induced fibrosis^3,4^. Reversal of pulmonary fibrosis by vaccinia immunotherapy requires creation of a lung-specific memory CD4^+^ Th1 cell^3^. Furthermore, our data show that vaccinia vaccination induced IFNγ producing TRM in the bleomycin-injured lungs whereas the unvaccinated fibrotic control lungs have a predominance of IL17^+^ TRM^3^. In this study, we demonstrate that IT adoptive transfer of CD49a^+^ TRMs are necessary and sufficient to reverse established murine fibrosis in the absence of vaccination.

The development of fibrosis is a complex process involving multiple cell types in the lung. Indeed, numerous studies, both mouse and human, have implicated not only fibroblasts, epithelial and immune cells but also a host of cytokines, chemokines and transcription factors as playing essential roles in the excess accumulation of ECM characteristic of lung fibrosis. We believe that these tissue-specific adoptively transferred TRMs interact with host immune cells to promote normal resolution of lung injury. Our bulk RNA seq data would also suggest that the CD49a^+^ TRM may be having broad effects on these processes that promote and maintain fibrosis. STRING analysis suggests that CD49a^+^ TRM upregulate distinct molecular pathways involved in collagen binding and signaling, extracellular matrix organization, metalloproteinases and growth factor and glycosaminoglycan binding. Thus, CD49a^+^ TRM reverse established fibrosis by promoting remodeling of the ECM, perhaps resetting the imbalance of injury to wound healing. Interestingly, the data demonstrating the paucity of CD49a^+^ cells in IPF lungs is consistent with our data that these cells are relatively absent in IPF lungs supporting a strategy of augmenting CD49a^+^ TRMs, via adoptive transfer as a means of promoting resolution of fibrosis and normal healing.

Over the last 30 years, much has been learned about lung fibrosis and the deadly human disease IPF. However, we still have not determined the cause, exact pathophysiology, or a cure for this invariably fatal disease. Like the recent advances using immunotherapy for cancer, we see a potential role for immunotherapy in IPF as well. Adoptive Cellular Therapy is emerging as a powerful means of treating certain cancers^36^. Our data provide the scientific rationale for the development of a novel paradigm shifting therapeutic approach to lung fibrosis in the form of immunotherapy of CD49a^+^ TRMs derived from a patient’s own T cells to be adoptively transferred into the lung to not only halt but reverse the progressive deadly fibrosis.

