## Supplemental Table 1 for "Adoptive transfer of CD49a^+^ Tissue resident memory cells reverses pulmonary fibrosis in mice"

| **Target** | **Target Species** | **Conjugate** | **Clone** | **Company** | **Product Number** |
| --- | --- | --- | --- | --- | --- |
| CD4 | Human | BV605 | OKT4 | Biolegend | 317437 |
| CD3 | Human | AF700 | UCHT1 | BD | 557917 |
| CD49a | Human | APC | TS2/7 | Biolegend | 328313 |
| TNFα | Human | BV421 | Mab11 | Biolegend | 502931 |
| IL17a | Human | AF488 | N49-653 | BD | 560489 |
| Granzyme B | Human | PE-CF594 | GB11 | BD | 562462 |
| IFNγ | Human | APC | 4S.B3 | Invitrogen | 53-7319-42 |
| CD49a | Human | Purified | TS2/7 | Biolegend | 328302 |
| CD103 | Mouse | PE/Dazzle 594 | 2E7 | Biolegend | 156909 |
| CD49a | Mouse | APC | HMa1 | Biolegend | 142606 |
| CD4 | Mouse | PE | RM4-4 | Biolegend | 116006 |
| CD4 | Mouse | BV650 | RM4-5 | Biolegend | 100546 |
| CD45 | Mouse | BV510 | 30-F11 | Biolegend | 103138 |
| IFNγ | Mouse | BV786 | XMG1.2 | Biolegend | 505838 |
| IL17a | Mouse | PE | TC11-18H10.1 | Biolegend | 506904 |

**Supplemental Table 1.**

Flow Cytometry Reagents

IHC Reagents

| Target | Target Species | Clone | Company | Product Number |
| --- | --- | --- | --- | --- |
| CD4 | Human | UMAB64 | Origene | UM800010 |
| CD49a | Human |  | Abcam | Ab181434 |
| CD103 | Human |  | Abcam | Ab129202 |
